# CZON-cutter: a CRISPR-Cas9 system with multiplexed organelle imaging in a simple unicellular alga

**DOI:** 10.1101/2021.05.23.445365

**Authors:** Naoto Tanaka, Yuko Mogi, Takayuki Fujiwara, Kannosuke Yabe, Yukiho Toyama, Tetsuya Higashiyama, Yamato Yoshida

## Abstract

The simple cellular structure of the unicellular alga *Cyanidioschyzon merolae* consists of one nucleus, one mitochondrion, one chloroplast, and one peroxisome per cell and offers unique advantages to investigate mechanisms of organellar proliferation and the cell cycle. Here, we describe an engineered clustered, regularly interspaced, short palindromic repeats (CRISPR)-associated protein 9 (Cas9) system, CZON-cutter, for simultaneous genome editing and organellar visualization. We engineered a *C. merolae* strain expressing a nuclear-localized Cas9-Venus nuclease to target editing at a locus defined by a single-guide RNA (sgRNA). We then successfully edited the algal genome and visualized the mitochondrion and peroxisome in transformants by fluorescent protein reporters with different excitation wavelengths. Fluorescent protein labeling of organelles in living transformants allows validation of phenotypes associated with organellar proliferation and the cell cycle, even when the edited gene is essential. Combined with the exceptional biological features of *C. merolae*, CZON-cutter will be instrumental for investigating cellular and organellar division in a high-throughput manner.

**Summary:** An engineered CRISPR-Cas9 system, named CZON-cutter, for simultaneous genome editing and fluorescent protein labeling of organelles in *Cyanidioschyzon merolae* can be used to validate intracellular function of a particular gene, even if it is essential.

## Introduction

Eukaryotic cells are composed of functionally specialized units represented by membrane-bound organelles, which respond to intercellular and intracellular signals for homeostasis and provide separated spaces in which to synthesize various biochemical molecules in varied metabolic pathways. Mitochondria and plastids are organelles surrounded by a double membrane that are thought to have evolved from endosymbiotic bacteria (Mereschkowsky, 1905; Gray, 1992; Martin and Kowallik, 1999). Because of their evolutionary origin, they carry the remnants of their own genome and only multiply via binary fission of pre-existing copies according to their own division cycles (Gillham et al., 1994; Kuroiwa et al., 1998; Suzuki et al., 1994; Kobayashi et al., 2011). Whether and how the cellular and nuclear, mitochondrial, and plastid division cycles cooperate in the cells of photosynthetic eukaryotes are still largely unknown.

To fill this gap in our knowledge, the field has recently focused on the simple unicellular alga *Cyanidioschyzon merolae*, commonly called “CZON” (Matsuzaki et al., 2004; Nozaki et al., 2007). *C. merolae* cells possess very few membrane-bound organelles: one nucleus, one mitochondrion, one plastid, one peroxisome, one Golgi body with two cisternae, a few vacuoles, and a simple shaped endoplasmic reticulum (ER). All organelles divide shortly before cell division and are then inherited by daughter cells (Imoto et al., 2010). Progression through the cell cycle can also be highly synchronized by entraining cultures to light/dark cycles (Suzuki et al., 1994). Together with its simple cell structure, these features provide *C. merolae* cells with unique advantages for investigating the molecular mechanisms related to organellar and cellular proliferation, as well as the cell cycle. Key factors for mitochondrial division (mitochondrial FtsZ, dynamin 1 [Dnm1], and MITOCHONDRION-DIVIDING RING1 [MDR1]) and for plastid division (plastid FtsZ, Dnm2, and PLASTID-DIVIDING RING1 [PDR1]) have been identified from a series of studies using *C. merolae* (Takahara et al., 2000; Nishida et al., 2003; Miyagishima et al., 2003; Yoshida et al., 2010, 2017). A subgroup of the kinesin super family has also been recently shown to be involved in chromosome segregation (Yoshida et al., 2013). Furthermore, a recent transcriptome analysis using synchronized *C. merolae* cultures identified 454 genes with an expression pattern driven by the cell cycle, of which 181 genes have no known function (Fujiwara et al., 2020). These unknown genes might contribute to the mechanism of organellar and cellular division.

Gene targeting approaches are indispensable and powerful tools to study the function of a gene. An optimized gene targeting technique has been established for *C. merolae* using homologous recombination (HR) and a positive selection method for uracil auxotrophy or chloramphenicol resistance (Minoda et al., 2004; Imamura et al., 2009; Fujiwara et al., 2013, 2017). However, HR-mediated deletion of a locus typically entails the assembly of a targeting construct consisting of 1-kb fragments specific for the target gene flanking the *URA5*.*3* selection marker before introduction into *C. merolae* cells (Fig. 1a). Furthermore, organellar morphology should be characterized by live microscopy observations using fluorescently labeled organelles in the resulting knockout cells to determine whether the targeted gene is involved in cellular and/or organellar division. The multi-step process of conventional gene targeting restricts investigation and is not efficient. Therefore, a simpler and higher-throughput method to inactivate gene function in *C. merolae* is needed.

**Figure 1.**
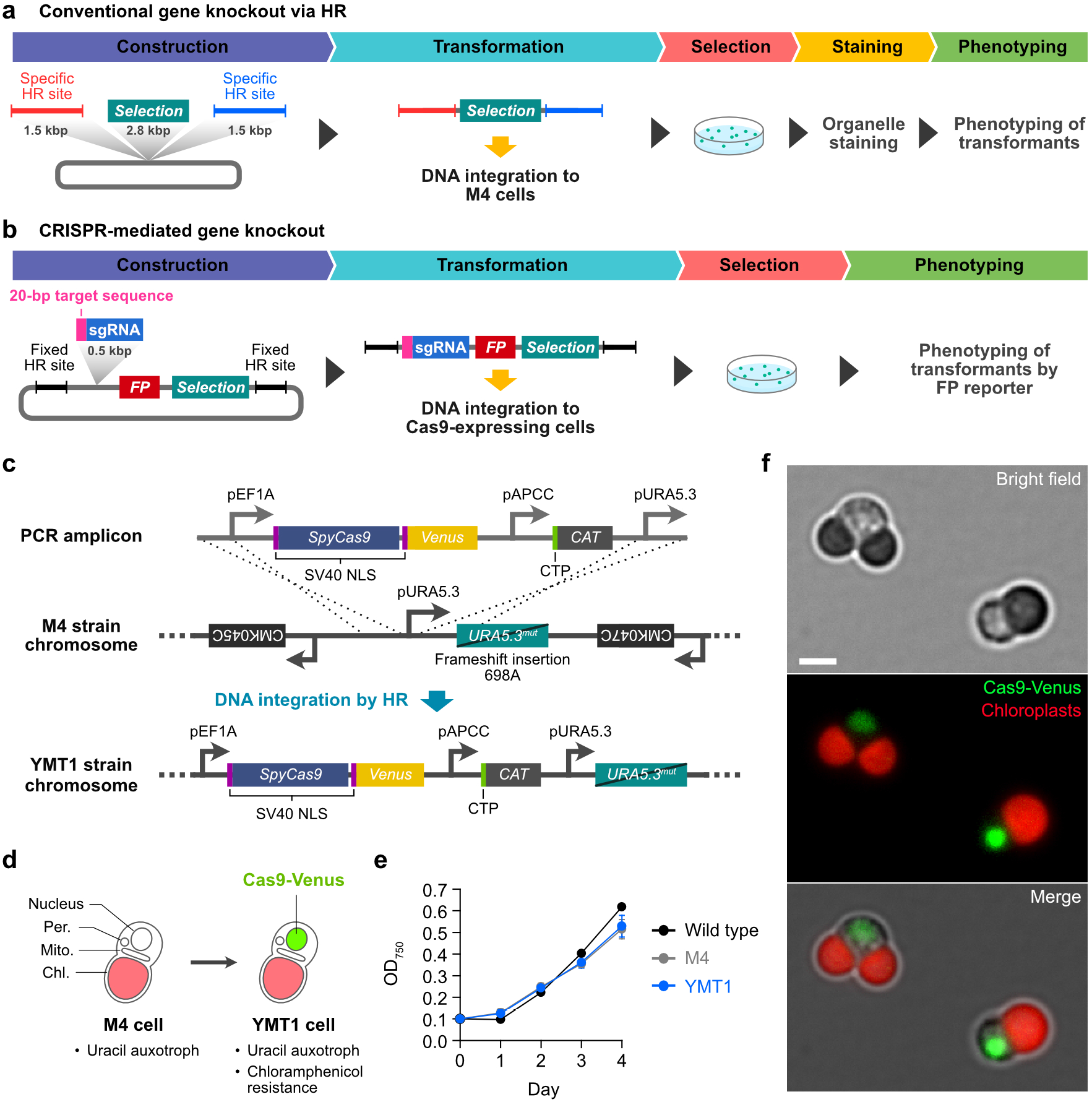
Development of the *Cyanidioschyzon merolae* YMT1 strain expressing Cas9-Venus. **(a)** Schematic depiction of a conventional gene targeting approach using homologous recombination in *C. merolae*. **(b)** Schematic depiction of a CRISPR-mediated gene targeting system with simultaneous organelle visualization. **(c)** Schematic diagram of insertion of the *Streptococcus pyogenes Cas9* gene fused with *Venus* (*Cas9-Venus*) and the selection marker *CAT* into the safe harbor site by homologous recombination. PCR amplicon, introduced linear DNA; M4 strain chromosome, genomic structure of the parental uracil-auxotrophic M4 strain, with the HR integration site. YMT1 strain chromosome, structure of the inserted *Cas9-Venus* and *CAT* cassettes in *C. merolae* strain YMT1. The *Cas9-Venus* cassette includes a 1-kb fragment of the *EF1A* promoter and 278 bp of the *UBQ3* 3’ untranslated region (UTR) for constitutive expression. Cas9-Venus was fused to two copies of a nuclear localization sequence (NLS) from SV40 T antigen to localize the Cas9-Venus fusion protein to the nucleus. *CAT* is driven by the constitutive *APCC* promoter with transcription termination by *β-Tubulin* 3’ UTR. The CAT coding sequence is preceded by a chloroplast transit peptide (CTP). The complete amino acid sequence of Cas9-Venus is given in Supplemental Note 1. **(d)** Schematic illustration of the *C. merolae* YMT1 strain. **(e)** Growth curves for the wild-type 10D, M4, and YMT1 strains. The 10D strain was cultured in 2× Allen’s medium. M4 and YMT1 strains were cultured in 2× Allen’s medium supplemented with uracil. **(f)** Imaging of a dividing cell (left) and a non-dividing cell (right) of *C. merolae* YMT1 strain. Green, Cas9-Venus fluorescence. Red, chlorophyll autofluorescence. Scale bar, 2 μm.

Targeted genome editing via clustered, regularly interspaced, short palindromic repeats (CRISPR) and CRISPR-associated protein 9 (Cas9) has revolutionized genetic analysis (Jiang and Doudna, 2017; Adli, 2018; Sander and Joung, 2014). The CRISPR-Cas9 system relies on the formation of a complex between Cas9 (most often the nuclease from *Streptococcus pyogenes*) and a single-guide RNA (sgRNA) whose sequence is complementary to 20 nucleotides (nt) of a sequence within the target gene, followed by a short DNA motif (protospacer-adjacent motif [PAM] sequence: NGG) (Nishimasu et al., 2014) that cleaves the genome at the target site. The resulting double-strand break can then be repaired by nonhomologous end-joining (NHEJ) or homology-directed repair (HDR) pathways in nearly all organisms. NHEJ-mediated repair frequently introduces insertion/deletion mutations (indels) of various lengths, which disrupt the open reading frame. When provided with exogenously supplied donor DNA templates, the HDR repair machinery can modify the genome by introducing specific point mutations or inserting desired sequences at the target site. The CRISPR-Cas9 system has therefore become a powerful tool for targeted genome editing in many established and emerging model organisms.

Here, to identify genes that are involved in organellar and cellular division, we describe an engineered CRISPR-Cas9 system for *C. merolae*, named CZON-cutter, that allows simultaneous site-selective genome editing and multiplexed organellar imaging. This CRISPR-based gene targeting system can be used in the engineered *C. merolae* strain YMT1, which expresses Cas9-Venus, upon transformation with plasmid DNA containing a designed sgRNA (Fig. 1b). Gene specificity can be achieved by altering the 20-nt target sequence of the sgRNA in the plasmid. Furthermore, the nucleus, single mitochondrion, and single peroxisome in transformed cells can be visualized by fluorescent protein reporters with different excitation wavelengths. The ability to image living transformants makes it possible to validate phenotypes associated with organellar morphology swiftly and accurately without chemical staining in the context of the cell cycle by time-lapse microscopy. The CZON-cutter platform thus holds great promise as an efficient, versatile, and high-throughput approach to investigate the biological function of any gene at organellar resolution.

## Results

### Engineering of the *Cyanidioschyzon merolae* YMT1 strain expressing Cas9-Venus and a universal plasmid DNA template containing a sgRNA and a mitochondrial reporter

To implement the CZON-cutter platform in *C. merolae*, we first fused the coding sequence for *Cas9* from *S. pyogenes* to two copies of a nuclear localization sequence (*NLS*) and the yellow fluorescent protein gene *Venus*. Because the resulting *NLS*-*Cas9-NLS-Venus* gene is driven by the constitutively expressed *ELONGATION FACTOR 1 ALPHA* (*EF1A*) promoter, Venus-tagged Cas9 (Cas9-Venus) should accumulate in the nucleus throughout the cell cycle. We then integrated a PCR amplicon consisting of the *Cas9-Venus* gene and a plastid-targeted *Chloramphenicol Acetyltransferase* (*CAT*) selection marker into a safe harbor site in the *C. merolae* uracil-auxotrophic M4 strain by HR (Fig. 1c). We selected transformants for resistance to chloramphenicol and named the resulting uracil-auxotrophic and chloramphenicol-resistant strain YMT1 (Fig. 1d). We confirmed that the YMT1 strain contains an inserted copy of the PCR amplicon and has a growth rate similar to that of the wild-type and M4 strains (Fig. 1e). Fluorescence microscopy of YMT1 cells demonstrated that Venus fluorescence is specifically detected in the nucleus of dividing and non-dividing cells (Fig. 1f). We thus concluded that Cas9-Venus accumulates in the nucleus throughout the cell cycle.

Next, to edit the genome and visualize mitochondria simultaneously, we generated another construct with a sgRNA and a fluorescent mitochondrial marker in the region downstream of *Cas9-Venus* by HR. For greater versatility, we constructed a universal plasmid DNA template, pGuide-mitoScarlet, containing two homologous regions and gene cassettes for the sgRNA, mitochondrion-targeted *mScarlet* gene (*mitoScarlet*), and the *URA5*.*3* selection marker (Fig. 2a). The sgRNA consisted of three segments: a 20-nt target-specific complementary region, a 76-nt scaffold region, and a 6-nt transcription termination signal (Fig. 2b). We placed the expression of the sgRNA under the control of a 593-bp promoter fragment from the *C. merolae* non-coding small nuclear RNA U6, which should be recognized by RNA polymerase III and thus highly expressed in *C. merolae* cells. The cassette encoding the mitochondrion-targeted fluorescent protein comprised the coding sequence for the red fluorescent protein mitoScarlet fused to a mitochondrial targeting sequence (MTS) driven by the constitutive *CPCC* promoter, followed by the *β-Tubulin* 3’ untranslated region (3’ UTR); this cassette was inserted into the plasmid downstream of *Venus* and upstream of *URA5*.*3*.

**Figure 2.**
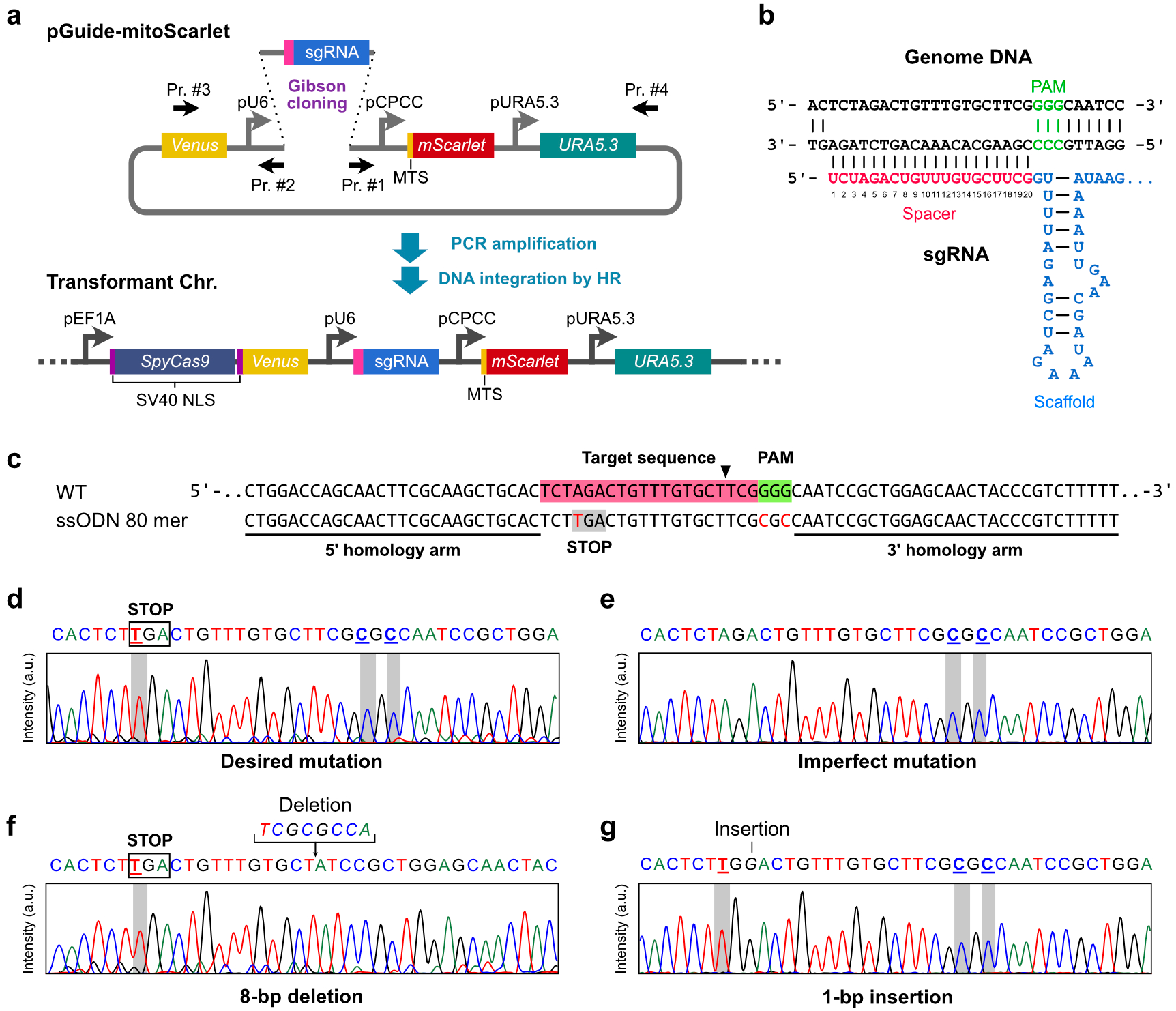
Design and implementation of CRISPR-based genome editing in *Cyanidioschyzon merolae*. **(a)** Site-specific insertion of gene cassettes containing a sgRNA, a mitochondrion-targeted red fluorescent protein (mitoScarlet), and the selection marker *URA5*.*3*. A synthetic single-guide RNA (sgRNA) is combined with a linearized plasmid amplified with primers #1 and #2 to generate pGuide-mitoScarlet (Fig. S3). A PCR amplicon obtained with primers #3 and #4 is then inserted into the region downstream of *Cas9-Venus* in the YMT1 strain by homologous recombination. MTS, mitochondrial targeting signal. **(b)** Design of the sgRNA and the target site for the *CRY* locus. Base-pairing nucleotides (20 bp) are shown in magenta, the Cas9-binding hairpin in blue, and the protospacer-adjacent motif (PAM) sequence in green. **(c)** Principle of Cas9-mediated precise genome editing of the *CRY* locus. The arrowhead indicates the putative Cas9 cleavage site. An 80-nt, single-stranded oligodeoxynucleotide (ssODN) donor template was designed to insert a stop codon and eliminate the PAM sequence in the *CRY* locus. The target sequence and the PAM are highlighted in magenta and green, respectively. The ssODN contains two 30-nt homology sequences (underlines). **(d–g)** Representative electropherograms of mutated *CRY* across 13 independent edited colonies. (**d**) All targeted nucleotides in *CRY* were accurately edited in 10 colonies. (**e**) Two of the three targeted nucleotides were edited in one colony. (**f**) An 8-bp deletion and (**g**) a 1-bp insertion were identified, in one colony each.

### CRISPR-based genome editing in *C. merolae* is performed by the HDR pathway

To validate the CRISPR-Cas9 gene editing system described above with the YMT1 strain and the pGuide-mitoScarlet plasmid, we selected the putative cryptochrome gene *CRY* (also called *PHR1*, accession number: CMO348C) as a target for genome editing. Cryptochromes in the fruit fly *Drosophila melanogaster* and other insects act as blue light circadian photoreceptors and in animals act as an integral component of the circadian machinery (Chaves et al., 2011; Öztürk et al., 2007), so we hypothesized that inactivation of *C. merolae CRY*, which clustered with animal and insect cryptochromes (Fig. S1), might disturb synchronization of cell division under light/dark cycles. Although the proteins encoded by the *C. merolae CRY* gene family exhibit photolyase activity to repair ultraviolet radiation–induced DNA damage, it remains unclear whether *C. merolae* CRYs behave as blue light photoreceptors (Asimgil and Kavakli, 2012). We designed the sgRNA target sequence against *CRY* with the web tool CRISPRdirect (https://crispr.dbcls.jp/) (Naito et al., 2015), by selecting a unique target sequence with a very low possibility of off-target effects. We then attempted to integrate a PCR amplicon, using the resulting pGuide-*CRY*_232-254_-mitoScarlet as template, into the genome of YMT1 cells by HR, but we failed to obtain a single positive genome-edited colony. This result suggested that *C. merolae* might not employ NHEJ to repair DNA double-strand breaks. In fact, the main components of the NHEJ pathway, *Ku70* and *Ku80* (Fell and Schild-Poulter, 2015), appear to be missing from the *C. merolae* genome. Therefore, we next tested Cas9-mediated genome editing via the HDR pathway. We used single-stranded oligodeoxynucleotides (ssODNs) as a donor template, consisting of two homology arms of 30 nt each flanking the 20-nt *CRY* target sequence, which contained three single-base substitutions, one introducing a stop codon and the other two eliminating the PAM sequence in the CRY target region (Fig. 2c). We then transfected the YMT1 strain with a PCR amplicon corresponding to pGuide-mitoScarlet-CRY with the ssODN to modify the target region of the *CRY* gene (Fig. 2d–2g). We obtained 16 colonies by the uracil-autotrophic selection and 13 positive clones by sequencing (∼11.1 transformants per 10^8^ cells). As each positive clone contains some kinds of nucleotide substitutions in the *CRY* target region, we verified that Cas9-Venus-mediated mutations occurred in the target region of the 13 positive clones to evaluate accuracy and variations of unexpected mutation in the genome editing. Ten clones (76.9%) harbored the desired mutations (Fig. 2d). The remaining three clones (23.1%) had imperfect and/or unexpected mutations at the *CRY* target site (Fig. 2e–2g). Furthermore, we detected red fluorescence in mitochondria from the accumulation of mScarlet and green fluorescence from Cas9-Venus in all 13 colonies by fluorescence microscopy (Fig. 3a). These results demonstrated successful genome editing and simultaneous visualization of the nucleus and mitochondrion in *C. merolae*; we named this CRISPR-based genome editing system CZON-cutter.

**Figure 3.**
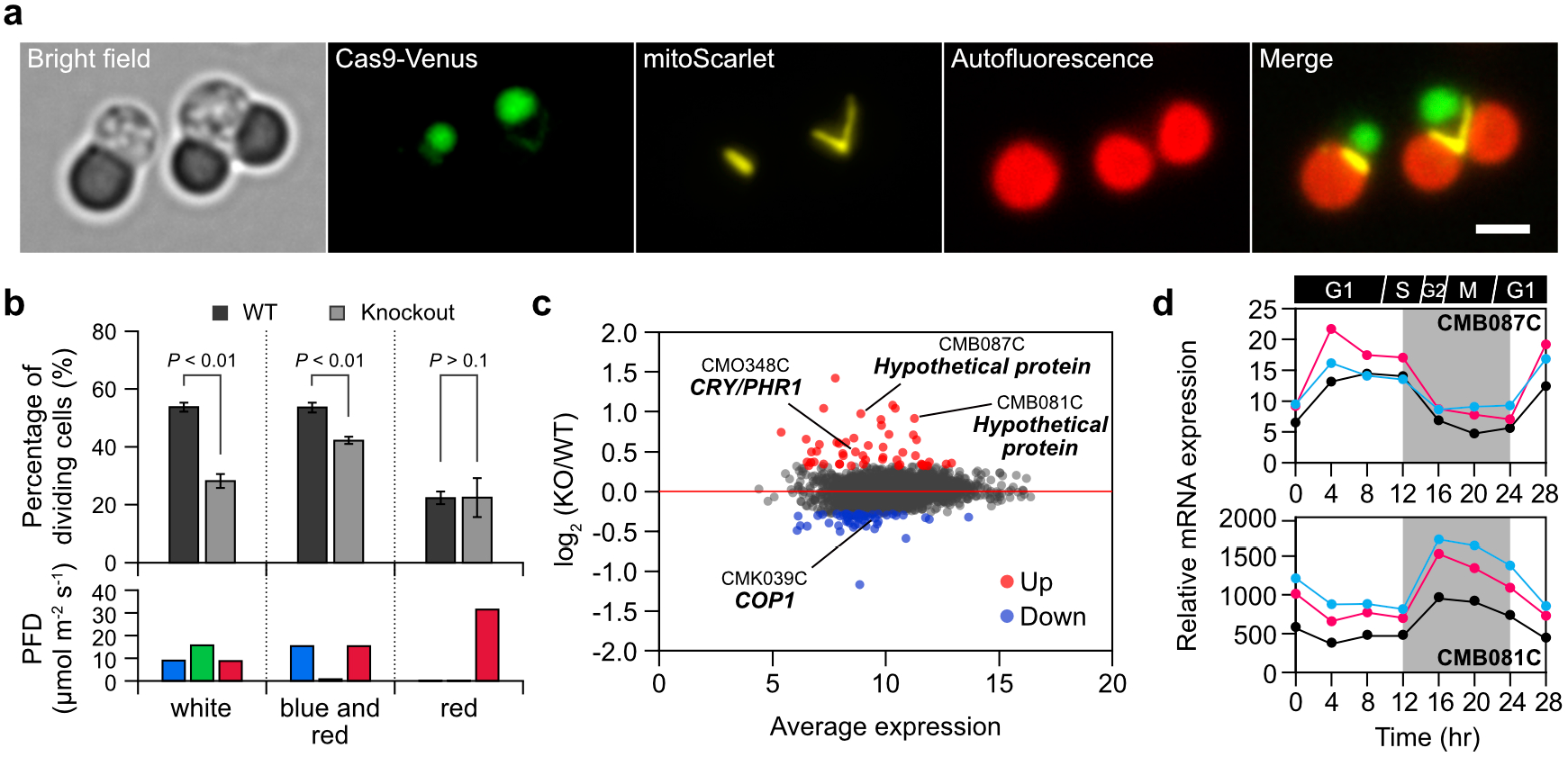
CRY-dependent circadian clock entrainment by blue light stimuli in *Cyanidioschyzon merolae*. **(a)** Imaging of a non-dividing cell (left) and a dividing cell (right) in the *CRY* knockout strain. Green, Cas9-Venus fluorescence; yellow, mitoScarlet fluorescence; red, chlorophyll autofluorescence. Scale bar, 2 μm. **(b)** Percentage of dividing cells in synchronized cultures for wild-type (WT) and *CRY* knockout strains under distinct light conditions. Total photon flux density (PFD) of blue, green, and red lights was kept at about 30 μmol m^−2^ s^−1^ in each light condition. Data are represented as means ± s.d. (*n* = 3 from individual experiments). *P* values are from two-sided Student’s *t*-test. Raw data are given in Table S1. **(c)** A MA plot of the fold-change [log_2_(KO/WT)] versus average expression of log_2_(TPM) from WT and *CRY* knockout (KO) samples. Expression values are given in Table S2. **(d)** Light-dependent expression changes of CMB087C and CMB081C. Black, WT strain; red, *CDKA* knockdown strain; blue, *RBR* knockout strain. Expression values in each strain were extracted from a time course (Fujiwara et al., 2020).

### Phenotype validation of the CRISPR-mediated *CRY* knockout strain by synchronization cultivation under distinct light conditions

To investigate whether *C. merolae CRY* affects cell-cycle progression, we next tested the synchronization of CRISPR-generated *CRY* loss-of-function mutants under light/dark cycles using one strain as a representative. For the purpose, we used not the uracil-auxotrophic YMT1 strain but the wild-type 10D strain for control experiments to compare in equal culture conditions using the non-uracil-supplemented medium. We cultured one *CRY* knockout strain and the wild-type 10D strain under light/dark cycles with white light (photon flux of 30 μmol m^−2^ s^−1^). Since a large percentage of *C. merolae* cells are in the dividing phase 38 hours after the initiation of synchronization (53.8% in this study), we compared the percentage of dividing cells in the knockout strain to that of the wild-type strain at this time point (Fig. 3b and Table S1). Fewer cells appeared to be dividing in the *CRY* knockout strain (28.2%, *p* value <0.01) relative to the wild-type strain, suggesting that inactivation of *CRY* may alter cell-cycle progression, possibly by disrupting circadian rhythms in *C. merolae*. Using the *CRY* knockout mutant strain, we next explored the effect of light quality on entrainment of the circadian clock in *C. merolae* (Fig. 3b and Table S1). We cultured the *CRY* knockout and the wild-type strain under light/dark cycles consisting of a combination of blue and red light irradiation. To maintain photosynthetic activity, we adjusted the total photon flux to provide approximately 30 μmol m^−2^ s^−1^ (blue: 15.4 μmol m^−2^ s^−1^, red: 15.4 μmol m^−2^ s^−1^). We observed that cells from the wild-type strain exhibit a comparable synchronization rate as that seen in white light, with a dividing rate of 53.6%. The percentage of dividing cells in the *CRY* knockout strain was significantly lower at 42.3% (*p* value <0.01), although it was higher than that of the *CRY* knockout strain grown in white light. As both the wild-type and the *CRY* knockout strains hardly proliferated under pure blue light, we could not confirm the synchronization properties of these strains under pure blue light condition. Therefore, we also evaluated whether *C. merolae* cell division was entrained by blue light by culturing cells under pure red light. Under these conditions, the percentages of dividing cells in the wild-type strain and the *CRY* knockout strain were identical at 22.4% and 22.5%, respectively, indicating that cell cycle of *C. merolae* is entrained by blue light irradiation. These findings suggest that *CRY* is involved in entrainment of the cell cycle by blue light in *C. merolae*. Because the cell cycle is gated by the circadian clock, these results suggest that *C. merolae* CRY may contribute to circadian entrainment.

### Identification of putative components in the regulation mechanism of the circadian rhythms in *C. merolae* by the genome-wide transcriptome using the *CRY* knockout strain

To explore the molecular links between cellular and organellar division and circadian rhythms in *C. merolae*, we compared gene expression profiles of the wild type and the *CRY* knockout mutant by transcriptome deep sequencing (RNA-seq) (Fig. 3c). We identified the 50 most highly upregulated genes and the 50 most strongly downregulated genes between the two strains (Table S2). *CRY* itself was among the upregulated genes, suggesting that an uncharacterized transcriptional activator may induce *CRY* expression to compensate for its loss. The expression level of the gene encoding the CRY regulatory protein COP1 was lower in the *CRY* mutant. COP1 was originally identified as a key factor repressing light-mediated development and growth in plants by degrading photomorphogenic transcription factors in the dark (Lau and Deng, 2012). In addition, CRYs antagonize COP1 activity to regulate circadian rhythms in mammals (Rizzini et al., 2019). As the loss of CRY function affected cell synchronization and the expression levels of *COP1, C. merolae* CRY may also play a pivotal role in the regulation of circadian rhythms in this alga. RNA-seq analysis indicated that 40% of the differentially expressed genes have unknown biological functions in the *C. merola*e genome database, hinting that some of these genes might act as nodes linking the circadian clock with the cellular and organellar division cycle. We independently validated the expression pattern of two of the most highly upregulated genes in our dataset, CMB087C and CMB081C, over a diurnal time course, with a peak in expression in the middle of the light period (CMB087C) or dark period (CMB081C), respectively (Fig. 3d). This diurnal pattern remained unchanged in knockout strains of either the *CYCLIN-DEPENDENT KINASE A* (*CDKA*) or *RETINOBLASTOMA-RELATED* gene (*Rb*). Thus, further exploration of the function of these genes may reveal the regulatory system underlying cellular/organellar division and circadian rhythms.

### Gene cassette knock-in for knocking out a target gene and fluorescent protein labeling of the peroxisome by CZON-cutter

We next used CZON-cutter to insert a sequence of interest into the genome (knock-in) while also knocking out a target gene and marking the single peroxisome with another fluorescent protein. We selected *ACTIN* as a target for gene knock-in. Although the *C. merolae* genome contains a single *ACTIN* gene (Takahashi et al., 1995; Matsuzaki et al., 2004), cytokinesis in *C. merolae* does not involve actin (Yagisawa et al., 2020). As actin plays a central role in cell division in contemporary eukaryotes, direct evidence of actin-free cell division in *C. merolae* would illustrate a new framework for cytokinesis in simpler eukaryotes. To generate a construct for CRISPR-mediated gene knock-in, we modified the sgRNA cloned into the pGuide-mitoScarlet plasmid to target *ACTIN* (Fig. 4a). We also constructed a plasmid that would act as template for HDR, pCer3-PTS1, in which the constitutive *APCC* promoter drives the expression of the gene encoding cyan fluorescence protein mCerulean3 fused to a peroxisome targeting signal (mCerulean3-PTS1, hereafter perCerulean3), followed by the *β-Tubulin* 3’ UTR. Target site specificity was provided by the PCR primers, which included 50-nt overhangs with homology to the target locus of interest. We transformed strain YMT1 with a pool of these two PCR products (one derived from pGuide-mitoScarlet and one from pCer3-PTS1) and obtained three uracil-autotrophic colonies expressing both mitoScarlet and perCerulean3 (∼2.6 transformants per 10^8^ cells). We chose one strain, the *ACTIN*^knockout^ + *perCerulean3*^knock-in^ strain, as a representative for further analysis and confirmed the presence of *perCerulean3* at the *ACTIN* locus (Fig. 4b; Fig. S2). We used fluorescence microscopy to image the single peroxisome, the nucleus, and the single mitochondrion in the same cell (Fig. 4c). We noticed no significant differences in the growth curves or cell shapes of the wild-type strain and the *ACTIN*^knockout^ + *perCerulean3*^knock-in^ strain over the cell cycle (Fig. 4d and 4e), and observation via fluorescence microscopy demonstrated that cells lacking actin undergoing cytokinesis successfully divide into two daughter cells, each with its own nucleus, mitochondrion, chloroplast, and peroxisome (Fig. 4e). We concluded that actin is not required for cell growth or proliferation in *C. merolae* and that the CZON-cutter platform can insert the *perCerulean3* cassette with 50-nt short homology arms into any locus of interest.

**Figure 4.**
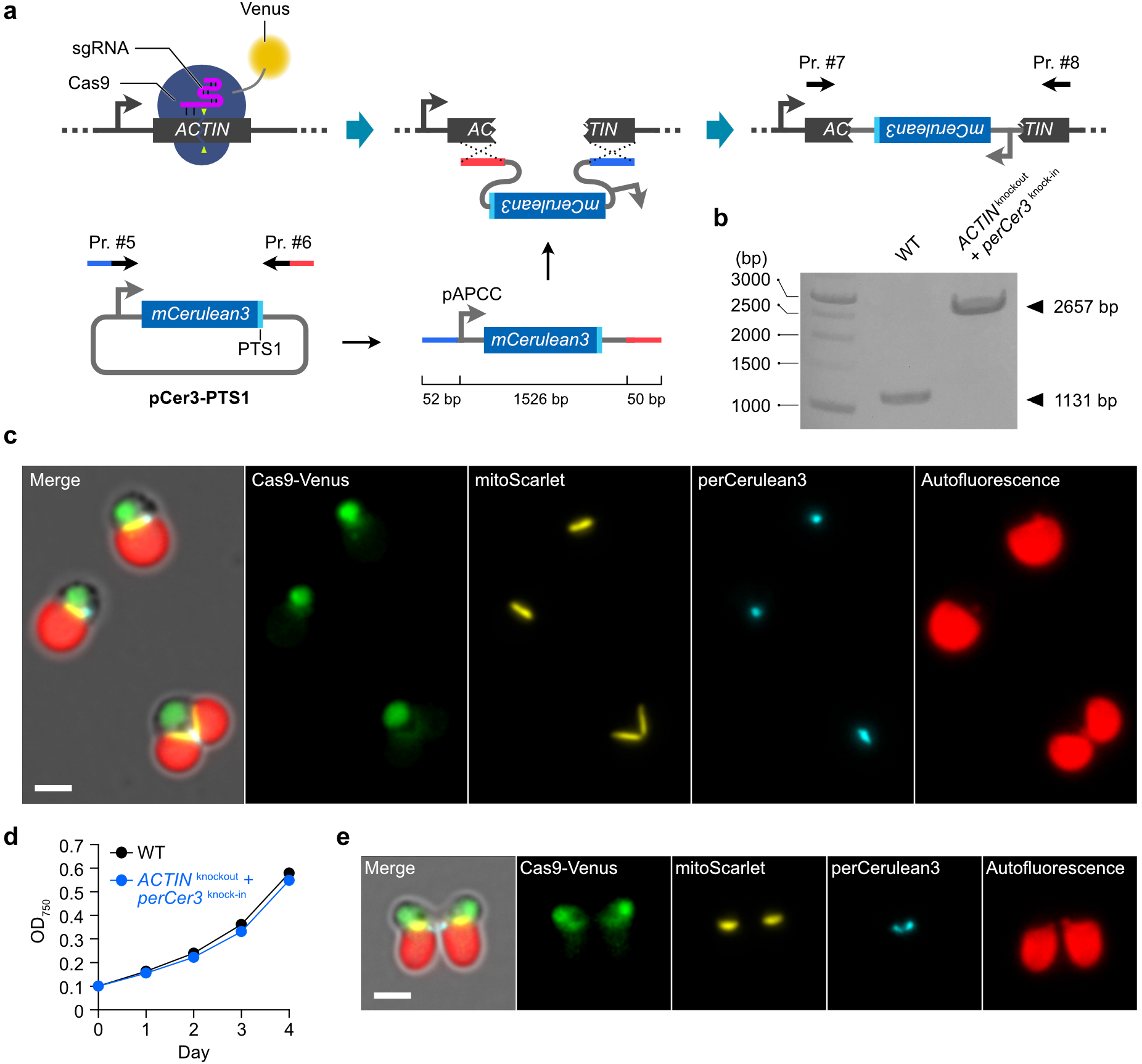
Site-specific knock-in of a cassette encoding *mCerulean3* fused to a peroxisomal targeting signal by CZON-cutter. **(a)** Principle behind knocking in a fluorescent reporter cassette at the *ACTIN* locus. The *mCerulean3* coding sequence with a peroxisomal targeting signal (*perCerulean3*) was amplified by PCR using primers #5 and #6. The PCR amplicon contains homology arms of ∼50 bp on either side of the cassette (which is 1,626 bp). PTS1, peroxisomal targeting signal 1. **(b)** Confirmation of the knock-in event at the *ACTIN* locus by PCR with primers #7 and #8 shown in (a). The wild-type (WT) strain was used as a negative control. **(c)** Representative images of a non-dividing cell (top and middle) and a dividing cell (bottom) of the *ACTIN*^knockout^ + *perCerulean3*^knock-in^ strain. Green, Cas9-Venus fluorescence; yellow, mitoScarlet fluorescence; blue, perCerulean3 fluorescence; red, chlorophyll autofluorescence. **(d)** Growth curves of the wild-type strain and the *ACTIN*^knockout^ + *perCerulean3*^knock-in^ strain. Data are represented as means ± s.d. (*n* = 3 from cell culture replicates). **(e)** Representative image of a single cell from the *ACTIN*^knockout^ + *perCerulean3*^knock-in^ strain at the cytokinetic abscission stage. Scale bars, 2 μm.

### Monitoring of living transformants to evaluate the effect of knocking out essential genes

Next, we assessed the applicability of the CZON-cutter platform to test the potential contribution of a given gene during organellar and cellular division, even when the gene is essential for growth and proliferation. For this purpose, we modified the sgRNA in the pGuide-mitoScarlet plasmid to target *MDR1* or *γ-Tubulin*. MDR1 participates in the assembly of the mitochondrion-dividing (MD) ring, the molecular machinery that severs the mitochondrion for mitochondrial proliferation (Yoshida et al., 2017), while γ-Tubulin plays a critical role in spindle assembly and chromosome segregation during M phase (Wiese and Zheng, 2006). We predicted that inactivation of MDR1 or γ-Tubulin would result in severe, abnormal mitochondrial division and cell division phenotypes. To evaluate the effect of knocking out these genes, we first investigated the morphology of transformants 2 days after transformation with PCR amplicons for the sgRNA targeting *MDR1* and the *perCerulean3* fluorescence reporter. We identified cells with fluorescence from mScarlet in the mitochondrion and mCerulean3 in the peroxisome, which comprised about 0.1% of all cells, indicative of our transformation efficiency (Fig. 5a). Since we detected fluorescence from both reporters (mitoScarlet for sgRNA and perCerulean3 for gene knock-in), we hypothesized that the *perCerulean3* cassette is integrated at the *MDR1* locus, thereby knocking it out. The morphological examination of transformants showed an obstruction of mitochondrial division and mitochondrion overgrowth. Furthermore, we also noticed nuclear division defects, peroxisomal division defects, and enlargement of cells. The fluorescence derived from Cas9-Venus allowed us to identify the nucleus in each cell; together with the presence of an overgrown mitochondrion, these observations suggested that the nuclear envelope is intact and that the transformant is arrested in prophase. The observed phenotype of mitochondrial division defects is very similar to that previously reported for antisense suppression of *MDR1* (Yoshida et al., 2017), suggesting that the gene knockout approach with CZON-cutter will be an effective method for the study of genes related to organellar and cellular division. Finally, we knocked out *γ-Tubulin* and characterized the resulting phenotypes with the same strategy. Two days after transformation, we identified cells exhibiting mitoScarlet and perCerulean3 fluorescence at a similar frequency (∼0.1%) as with *MDR1* (Fig. 5b). Although we detected strong fluorescence for Cas9-Venus in the putative nuclear region, we also observed some fluorescence in the cytosol, indicating that the nuclear envelope is partially disassembled and that the transformant is arrested in prometaphase or metaphase. In addition, the chloroplast and mitochondrion had already undergone division, while we observed a single peroxisome in one of the two dividing cells. The results indicated that the division of each organelle—chloroplast, mitochondrion, and peroxisome—occurs at different times during the cell cycle (Fig. 5c and 5d). The CZON-cutter platform will be instrumental in exploring the function of genes involved in organellar and cellular division, even if the gene is essential.

**Figure 5.**
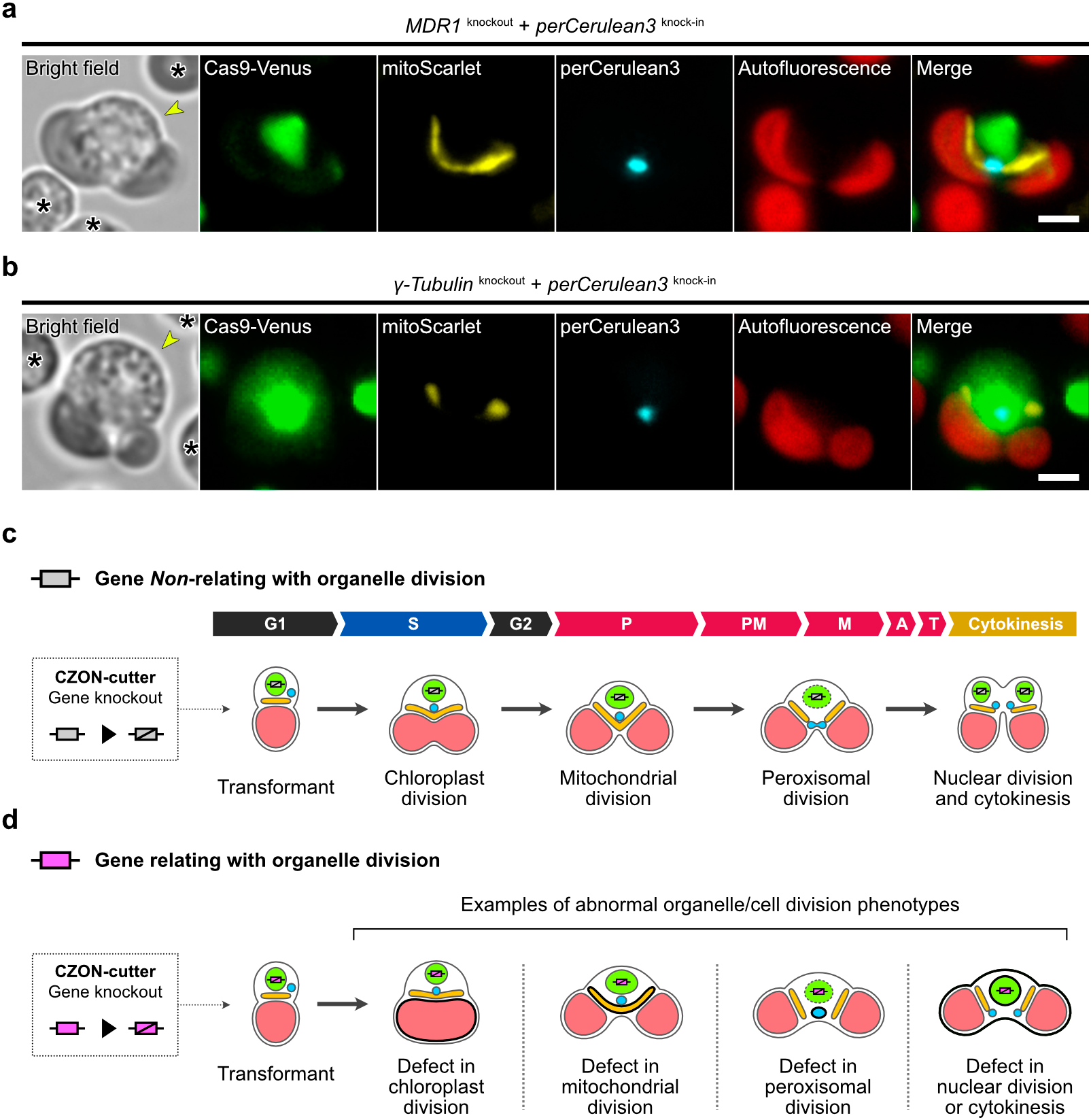
Essential genes for organellar and cellular division can be targeted by CZON-cutter. **(a and b)** Representative images of a *MDR1* knockout cell and a *γ-Tubulin* knockout cell. The cells were imaged 2 days after transformation. As these are living cells, they floated in the mounting medium and shifted slightly during imaging; the transformed cell (yellow arrowhead) is shown in the center of the field, while non-transformed cells are indicated with asterisks. Each gene was knocked out by inserting the *perCerulean3* cassette at the target locus. Scale bars, 2 μm. **(c and d)** Schematic illustrations of putative morphological phenotypes of transformed cells via CZON-cutter. When a target gene does not affect organellar and/or cellular division, transformed cells undergo normal organellar divisions and cytokinesis, as shown here. However, when a target gene is involved in division, the resulting transformed cells may become arrested at a particular cell division phase with abnormal morphology specific for each organellar division defect.

## Discussion

### CRISPR-mediated genome editing with multiplexed organelle visualization in *C. merolae*

The unique cell structure and availability of the complete genome sequence of *C. merolae* offers a new avenue to study intracellular mechanisms of eukaryotic cells. Here, we describe an efficient and accurate gene targeting method to take advantage of the unique features of this unicellular alga. Conventional HR techniques in *C. merolae* involve the introduction of a linear DNA fragment consisting of a gene cassette and selection marker flanked by over 1 kb of sequence from the locus of interest on either side (Fujiwara et al., 2017). The generation of a transformation construct therefore entails multiple steps, and the resulting plasmid is rather complex (Fig. 1a). Although *C. merolae* boasts the smallest genome of all photosynthetic eukaryotes, cumbersome gene targeting has hindered genome-wide analyses of gene function in this alga. Here, we established the CRISPR-mediated genome-editing platform CZON-cutter, which can disrupt or modify any gene in the genome with the added benefit of multiplexed visualization of organelles in a simple procedure (Fig. 1b; Figs. S3 and S4). The engineered strain used here, YMT1, showed no abnormal growth phenotype (Fig. 1e). Given the small size of the *C. merolae* genome, potential off-target effects by CZON-cutter should be minimal. Furthermore, multiplexed visualization of organelles by high-brightness fluorescence proteins allows the prompt characterization of the phenotypes associated with a gene knockout in terms of organellar division and morphology.

### The CZON-cutter platform provides an efficient, versatile, and high-throughput approach to investigate the biological function of any gene at organellar resolution

Mitochondria and chloroplasts proliferate via binary division of preexisting organelles by using specialized ring-shaped complexes called mitochondrial- and plastid-division machineries (Yoshida and Mogi, 2019; Yoshida et al., 2010, 2017). Peroxisomes divide by using a homologous division machinery (Imoto et al., 2013). However, the individual components and the underlying regulatory mechanisms of these division machineries are largely unknown. A recent transcriptome study identified 181 uncharacterized genes that are specifically expressed during mitochondrial and chloroplast division (Fujiwara et al., 2020), which may be candidate genes for exploring the mechanisms behind organellar division.

In this study, we generated a knockout strain for *CRY* with CZON-cutter, which revealed that CRY may be a blue light photoreceptor with critical roles in the regulation of cell-cycle progression in *C. merolae*. CZON-cutter also paves the way for studies of the physiological and molecular mechanisms of the cell cycle. Furthermore, the successful knockout of *MDR1* or *γ-Tubulin* also demonstrated that CZON-cutter makes it possible to evaluate the function of essential genes (Fig. 5). In addition, our results hint at the links between organellar division and cellular division. A defect in mitochondrial division caused by genome editing of *MDR1* arrested cell-cycle progression at prophase, indicating that a mitochondrial division may participate in a prophase–prometaphase checkpoint. Another notable illustration of the power of CZON-cutter is evident in the gene knockout of *γ-Tubulin*, which not only blocked chromosome segregation but also exhibited a peroxisomal division defect. These results indicated that chloroplast, mitochondrial, and peroxisomal divisions are executed in a specific sequence during the cell cycle and that completion of each organellar division is likely to serve as a checkpoint in the cell cycle. Further studies will aim to explore the molecular mechanisms of this highly interdependent system between division of organelles and the cell. In conclusion, CZON-cutter enables the systematic analysis of gene function by genome editing and multiplexed organellar visualization. The originality and versatility of CZON-cutter will help accelerate gene discovery to reveal biological principles of the cell.

## Materials and methods

### *C. merolae* YMT1 strain

To create a *C. merolae* strain constitutively accumulating nuclear-localized Cas9 fused to Venus, a construct containing *Cas9-Venus* and chloroplast-targeted *CAT* was assembled as follows. The *EF1A* promoter (*EF1Ap*, 1 kb) was PCR-amplified using genomic DNA from *C. merolae* strain 10D as template. A version of *S. pyogenes Cas9*, codon-optimized for mammalian expression and fused to three copies of the FLAG tag and two copies of a nuclear localization signal from simian virus 40 (SV40), was PCR-amplified from plasmid pKIR1.1 (Addgene). *Venus* codon-optimized for expression in *C. merolae* (Nagai et al., 2002) and fused to three copies of the HA tag, the *UBIQUITIN 3* (*UBQ3*) 3’ UTR, *APCC* promoter, *CAT* fused to a chloroplast transit peptide (CTP) derived from APCC, and *β-Tubulin* 3’ UTR were amplified by PCR using plasmid pVenus-CAT (gift from T. Fujiwara) as template. The resulting PCR amplicons were cloned into a linear vector amplified by PCR using p*NIR*p::sfGFP (gift from T. Fujiwara) (Fujiwara et al., 2015) as template. The resulting construct, pCas9-Venus-CAT, contained the gene cassettes *Cas9-Venus* and *CTP-CAT* approximately 0.9 kb upstream of *URA5*.*3*. For the construction of pCas9-Venus-CAT, PCR amplification and assembly of DNA fragments were performed with Platinum SuperFi II DNA polymerase (Thermo Fisher) and the NEBuilder HiFi DNA assembly cloning kit (New England Biolabs), respectively. Finally, using pCas9-Venus-CAT as template, a PCR amplicon was amplified that contained ∼1,400 bp of sequence upstream of *URA5*.*3* (2,300 bp to 898 bp), the *EF1A* promoter, *Cas9-Venus*, the *UBQ3* 3’ UTR, the *APCC* promoter, *CTP-CAT*, the *β-Tubulin* 3’ UTR, and ∼900 bp of sequence upstream of *URA5*.*3* (897 to 1 bp). The PCR amplicon was introduced into the uracil-auxotrophic mutant M4 cells by polyethylene glycol (PEG)-mediated transformation as described by Ohnuma et al. (2008), Imamura et al. (2008), and Fujiwara et al. (2015). Chloramphenicol-resistant transformants were selected by cultivation in chloramphenicol-containing medium at 150 μg mL^−1^ for 15 days as described by Fujiwara et al. (2017). After chloramphenicol selection, transformants were spread on modified Allen’s (MA) medium supplemented with uracil (0.5 mg mL^−1^) to isolate single colonies. Accumulation of Cas9-Venus was examined by fluorescence microscopy, and the presence of the introduced DNA was confirmed by direct colony PCR and sequencing. The full amino acid sequence of the Cas9-Venus is given in Supplemental Note 1.

### Construction of sgRNA expression vectors

To generate the sgRNA expression vector, the *C. merolae* U6 promoter and a sgRNA scaffold with a termination signal were synthesized. The synthetic DNA fragments were assembled with a linear vector DNA containing 2,300 bp of *URA5*.*3* upstream sequence and the *URA5*.*3* open reading frame, PCR-amplified using p*NIR*p::sfGFP (Fujiwara et al., 2015) as a template. Then, the *UBQ3* 3’ UTR, the *CPCC* promoter, the mitochondrial transit signal (MTS) derived from mitochondrial *EF-Tu, mScarlet* (Bindels et al., 2016) codon-optimized for *C. merolae* expression, and the *β-Tubulin* 3’ UTR were introduced into the vector. Finally, the *URA5*.*3* upstream sequence (from 2,300 to 898 bp) was replaced by *Venus* and the *UBQ3* 3’ UTR to create a HR site to introduce constructs into the YMT1 strain. Potential Cas9 target sites in the *CRY, ACTIN, MDR1*, and *γ-Tubulin* loci were searched using the CRISPRdirect web server (http://crispr.dbcls.jp/) using the genome of *C. merolae* ASM9120v1 (Table. S3). To create the sgRNA expression vector targeting *CRY* (pGuide-*CRY*_232-254_-mitoScarlet), a 497-bp synthesized DNA fragment containing sgRNA spacer sequence targeting nucleotides 232–254 of the *CRY* locus was assembled with a linear vector amplified by PCR with primer set #1 and #2 using pGuide-mitoScarlet as the template. Similarly, the sgRNA expression vectors targeting *ACTIN* (pGuide-*ACTIN*_239-261_-mitoScarlet) and *MDR1* (pGuide-*MDR1*_254-276_-mitoScarlet) were constructed from synthesized DNA fragments containing a sgRNA spacer sequence targeting the *ACTIN* or *MDR1* locus, respectively. The complete nucleotide sequences of synthetic DNAs for sgRNAs targeting the *CRY, ACTIN*, and *MDR1* loci are given in Table S4 as #16 to #18. To create the sgRNA expression vector targeting *γ-Tubulin* (pGuide-*γTub*_863-885_-mitoScarlet), a 62-bp ssODN (#9) containing a sgRNA spacer sequence targeting nucleotides 863–885 bp of the *γ-Tubulin* locus was assembled with a linear vector amplified by PCR with primer sets #10 and #11 using pGuide-mitoScarlet as template. One guanine nucleobase was added at the 5’ side of the target sequence for *CRY* and *γ-Tubulin* to stimulate transcription of sgRNA *in vivo*.

A plasmid containing peroxisome-targeted *mCerulean3* codon-optimized for *C. merolae* expression was constructed as follows. *mCerulean3* (Markwardt et al., 2011) fused to peroxisomal targeting signal 1 (PTS1) was amplified by PCR with primer sets #19 and #20 using the above plasmid as template to generate peroxisome-targeted *mCerulean3* (*perCerulean3*). The *APCC* promoter, *perCerulean3*, the *β-Tubulin* 3’ UTR, and the pUC57 vector were amplified by PCR using primer sets #19 to #26 and assembled into a new vector named pCer3-PTS1. The full nucleotide sequence of *perCerulean3* is given in Supplemental Note 2. To carry out knock-in experiments, the *perCerulean3* cassette was amplified by PCR using pCer3-PTS1 as template with primer sets #5 and #6 for *ACTIN*, #12 and #13 for *MDR1*, and #14 and #15 for *γ-Tubulin*.

For all plasmids, PCR amplification and assembly of DNA fragments were performed using Platinum SuperFi II DNA polymerase (Thermo Fisher) and the NEBuilder HiFi DNA assembly cloning kit (New England Biolabs), respectively.

### Genome editing experiments

For genome editing of the *CRY* locus, a PCR amplicon was amplified from pGuide-*CRY*_232-254_-mitoScarlet as template and contained the sgRNA targeting nucleotides 232–254 of the *CRY* locus, mitochondrion-targeted red fluorescence protein mScarlet (*mitoScarlet*), and the selection marker *URA5*.*3*, flanked by HR sites. The PCR amplicon was then mixed with an 80-nt ssODN consisting of a 20-nt target sequence and two 30-nt flanking upstream and downstream sequences of the target sequence. The mixture was then transformed into YMT1 cells as described by Fujiwara et al. (2015). For gene knock-in at the target locus, PCR amplicons were amplified with primer sets #3 and #4 from pGuide-*ACTIN*_239-261_-mitoScarlet, pGuide-*MDR1*_254-276_-mitoScarlet, or pGuide-*γTub*_863-885_-mitoScarlet as templates; each PCR product consisted of the respective sgRNA, *mitoScarlet*, and *URA5*.*3* flanked with HR sites. In addition, the *perCerulean3* cassette was PCR-amplified with short homology arms with primer sets #5 and #6 (for *ACTIN*), #12 and #13 (for *MDR1*), or #14 and #15 (for *γ-Tubulin*) using pCer3-PTS1 as template. The resulting PCR amplicons (from pGuide and pCer3-PTS1) were then mixed and introduced into YMT1 cells as described by Fujiwara et al. (2015).

### Fluorescence microscopy

Fluorescence observations were conducted on an Olympus IX83 inverted microscope with a 1.45 NA, 100× oil immersion objective. Illumination was provided by using a fluorescence light source U-HGLGPS (Olympus), and the samples were observed through excitation filters [490-500HQ (Olympus) for Venus, FF01-549/12-25 (Semrock) for mScarlet, FF01-427/10-25 (Semrock) for mCerulean3 and FF01-405/10-25 (Semrock) for chloroplasts], custom dichroic mirrors [Di03-R514-t1-25×36 (Semrock) for Venus, Di03-R561-t1-25×36 (Semrock) for mScarlet, FF458-Di02-25×36 (Semrock) for mCerulean3 and T455lp (Chroma) for chloroplasts], emission filters [FF02-531/22-25 (Semrock) for Venus, FF02-585/29-25 for mScarlet (Semrock), FF01-474/27-25 for mCerulean3, and FF02-617/73-25 for chloroplasts (Semrock)]. Images were acquired with a Zyla 4.2 sCMOS camera (Andor), controlled by MetaMorph software (Molecular Devices). The effective pixel size was 65.2 nm.

### Cell cultures and synchronization cultivation

*C. merolae* 10D strain cells (NIES-3377) were used as the wild type in this study. Wild-type cells and CRISPR-generated mutants were maintained on 2× Allen’s medium. The uracil-auxotrophic M4 strain and the uracil-auxotrophic/chloramphenicol-resistant YMT1 strain were maintained on MA2 medium supplemented with uracil (0.5 mg mL^−1^) and 5-fluoroorotic acid monohydrate (0.8 mg mL^−1^). All strains were cultured in flasks with agitation at 120 rpm under continuous white light (22 μmol m^−2^ s^−1^) at 38°C.

For synchronization of cultivation, cells were subcultured to <1 × 10^7^ cells mL^−1^ in a 100-mL flask and bubbled with filtered clean and humid air through a tube connected to an aquarium pump. Cells were then incubated under a 12-hour-light/12-hour-dark cycle at 42°C in a SLI-700 incubator (EYELA). Blue and red light irradiation were supplied using 470 nm ± 20 nm LED lights ISL-150×150-HBB (CCS Inc.) and 660 nm ± 20 nm LED lights ISL-150×150-RR (CCS Inc.), respectively.

### Phylogenetic analysis

The accession numbers used were as follows: *Cyanidioschyzon merolae* CRY/PHR1 (XP_005537706), *C. merolae* PHR2 (BAM80259), *C. merolae* PHR3 (BAM80957), *C. merolae* PHR4 (BAM79915), *C. merolae* PHR5 (BAM78760), *C. merolae* PHR7 (BAM82280), *Arabidopsis thaliana CRY1* (AEE82696), *A. thaliana CRY2* (AEE27692), *A. thaliana CRY-DASH* (Q84KJ5), *Homo sapiens CRY1* (NP_004066), *H. sapiens CRY2* (Q49AN0), *Mus musculus CRY1* (AAD39548), *M. musculus CRY2* (AAD46561), *Gallus gallus* CRY1 (AAK61385), *G. gallus* CRY2 (AAK61386), *G. gallus* CRY4 (NP_001034685), *Drosophila melanogaster* CRY (NP_732407), *D. melanogaster* 6-4 photolyase (BAA12067), *Escherichia coli* DNA photolyase (WP_062883603), and *Synechocystis* sp. PCC 6803 DNA photolyase (Q55081).

The phylogenic tree was generated with MEGA X (Kumar et al., 2018). Amino acid sequences were aligned with the Clustal W algorithm with default settings in MEGA X using the Maximum Likelihood method with the LG+G+I model. The local probability of each branch was calculated using the Neighbor-Joining method with 1,000 replications.

### RNA sequencing and analysis

Total RNA was purified from cells using the Trizol/RNeasy hybrid protocol (Trizol, Life Technologies; RNeasy Mini Kit, Qiagen). Polyadenylated [poly(A)] RNA was then purified with the NEBNext Poly(A) mRNA Magnetic Isolation Module (NEB, E7490). Sequencing libraries were constructed with the NEBNext Ultra II Directional RNA Library Prep kit for Illumina (NEB, E7760), amplified with custom oligonucleotides, and sequenced as 150-bp paired-end reads on an Illumina NovaSeq sequencer at GENEWIZ Inc. NovaSeq paired-end reads were mapped to the *C. merolae* genome (ASM9120v1) using bowtie2 and counted by featureCounts (Langmead and Salzberg, 2012; Liao et al., 2014). Genes with over 100 reads were selected and their expression normalized as transcripts per kb million (TPM). We identified the top up-and downregulated genes between the wild-type and the *CRY1* knockout strains from the log_2_ (fold-change) of TPM-normalized read counts. The genes are listed in Table S2.

### Online supplemental material

Fig. S1 shows a phylogenetic tree of cryptochromes and photolyases. Fig. S2 shows DNA sequence at the insertion site of *perCerulean3* knocked in to *ACTIN* by CZON-cutter. Fig. S3 shows construction of the pGuide-mitoScarlet plasmid targeting a target locus. Fig. S4 shows schematic flowchart of CZON-cutter. Table S1 describes summary of data sets for synchronization of the wild-type (WT) and *CRY* knockout (KO) strains. Table S2 describes top 50 upregulated and downregulated genes between the *CRY* knockout strain and the wild-type strain. Table S3 describes evaluation of CRISPR-Cas9 target sites for *CRY, ACTIN, MDR1*, and *γ-Tubulin* by the CRISPRdirect online tool. Table S4 shows primers and synthetic DNA fragments used in this study. Supplemental note 1 describes complete amino acid sequence of Cas9-Venus. Supplemental note 2 describes complete nucleotide sequence of *Cyanidioschyzon merolae* codon-optimized *mCerulean3* with the sequence for a peroxisomal targeting signal 1 (PTS1).

## Acknowledgments

This work was supported by PRESTO from the Japan Science and Technology Agency (JPMJPR20EE to Y.Y.); the Human Frontier Science Program Career Development Award (no. CDA00049/2018-C to Y.Y.); Japan Society for the Promotion of Science KAKENHI (no. JP18K06325 to Y.Y. and 18K06300 to T.F.); the Sumitomo Foundation (no. 180705 to Y.Y.); the Institution for Fermentation, Osaka (L-2020-2-008 to Y.Y.); the Grant-in-Aid for Scientific Research on Innovative Areas (no. 16H06465 to T.H. and Y.Y.); and CREST from the Japan Science and Technology Agency (JPMJCR20E5 to T.H.).

The authors declare no competing or financial interests.

## Author contributions

N.T., Y.M., T.F., and Y.Y designed and generated the YMT1 strain. N.T., Y.M., K.Y., and Y.Y. performed transformation experiments. N.T., Y.M., T.F., Y.T. and Y.Y. designed and constructed the sgRNA vector pGuide-mitoScarlet and a gene-cassette knock-in vector pCer3-PTS1. Y.M., K.Y., and Y.Y. performed analysis of CRISPR-mediated mutants. Y.M. and Y.Y. performed synchronization cultivation and comparisons of synchronization properties. N.T., Y.M., and Y.Y. collected microscopy images. N.T., Y.M., and Y.Y. performed RNA-seq experiments and analyzed data. N.T., Y.M., T.H. and Y.Y. wrote the manuscript with help from all authors. Y.Y. directed and supervised all research.

## Notes

### Competing Interest Statement

The authors have declared no competing interest.

## References

Adli, M. 2018. The CRISPR tool kit for genome editing and beyond. Nat. Commun. 9:1911. doi:10.1038/s41467-018-04252-2.

Asimgil, H., and I.H. Kavakli. 2012. Purification and characterization of five members of photolyase/cryptochrome family from Cyanidioschyzon merolae. Plant Sci. 185–186:190–198. doi:10.1016/j.plantsci.2011.10.005.

Bindels, D.S., L. Haarbosch, L. Van Weeren, M. Postma, K.E. Wiese, M. Mastop, S. Aumonier, G. Gotthard, A. Royant, M.A. Hink, and T.W.J. Gadella. 2016. mScarlet: a bright monomeric red fluorescent protein for cellular imaging. Nat. Methods. 14:53–56. doi:10.1038/nmeth.4074.

Chaves, I., R. Pokorny, M. Byrdin, N. Hoang, T. Ritz, K. Brettel, L.-O. Essen, G.T.J. van der Horst, A. Batschauer, and M. Ahmad. 2011. The Cryptochromes: Blue Light Photoreceptors in Plants and Animals. Annu. Rev. Plant Biol. 62:335–364. doi:10.1146/annurev-arplant-042110-103759.

Fell, V.L., and C. Schild-Poulter. 2015. The Ku heterodimer: Function in DNA repair and beyond. Mutat. Res. - Rev. Mutat. Res. 763:15–29. doi:10.1016/j.mrrev.2014.06.002.

Fujiwara, T., S. Hirooka, R. Ohbayashi, R. Onuma, and S.Y. Miyagishima. 2020. Relationship between cell cycle and diel transcriptomic changes in metabolism in a unicellular red alga1[OPEN]. Plant Physiol. 183:1484–1501. doi:10.1104/pp.20.00469.

Fujiwara, T., Y. Kanesaki, S. Hirooka, A. Era, N. Sumiya, H. Yoshikawa, K. Tanaka, and S.-Y. Miyagishima. 2015. A nitrogen source-dependent inducible and repressible gene expression system in the red alga Cyanidioschyzon merolae. Front. Plant Sci. 6:1–10. doi:10.3389/fpls.2015.00657.

Fujiwara, T., M. Ohnuma, T. Kuroiwa, R. Ohbayashi, S. Hirooka, and S.-Y. Miyagishima. 2017. Development of a Double Nuclear Gene-Targeting Method by Two-Step Transformation Based on a Newly Established Chloramphenicol-Selection System in the Red Alga Cyanidioschyzon merolae. Front. Plant Sci. 8:1–10. doi:10.3389/fpls.2017.00343.

Fujiwara, T., K. Tanaka, T. Kuroiwa, and T. Hirano. 2013. Spatiotemporal dynamics of condensins I and II: Evolutionary insights from the primitive red alga Cyanidioschyzon merolae. Mol. Biol. Cell. 24:2515–2527. doi:10.1091/mbc.E13-04-0208.

Gillham, N.W., J.E. Boynton, and C.R. Hauser. 1994. Translational regulation of gene expression in chloroplasts and mitochondria. Annu. Rev. Genet. 28:71–93. doi:10.1146/annurev.ge.28.120194.000443.

Gray, M.W.W. 1992. The Endosymbiont Hypothesis Revisited. Int. Rev. Cytol. 141:233–357. doi:10.1016/S0074-7696(08)62068-9.

Imamura, S., Y. Kanesaki, M. Ohnuma, T. Inouye, Y. Sekine, T. Fujiwara, T. Kuroiwa, and K. Tanaka. 2009. R2R3-type MYB transcription factor, CmMYB1, is a central nitrogen assimilation regulator in Cyanidioschyzon merolae. Proc. Natl. Acad. Sci. 106:14180–14180. doi:10.1073/pnas.0908318106.

Imoto, Y., T. Fujiwara, Y. Yoshida, H. Kuroiwa, S. Maruyama, and T. Kuroiwa. 2010. Division of cell nuclei, mitochondria, plastids, and microbodies mediated by mitotic spindle poles in the primitive red alga Cyanidioschyzon merolae. Protoplasma. 241:63–74. doi:10.1007/s00709-010-0107-y.

Imoto, Y., H. Kuroiwa, Y. Yoshida, M. Ohnuma, T. Fujiwara, M. Yoshida, K. Nishida, F. Yagisawa, S. Hirooka, S. Miyagishima, O. Misumi, S. Kawano, and T. Kuroiwa. 2013. Single-membrane-bounded peroxisome division revealed by isolation of dynamin-based machinery. Proc Natl Acad Sci U S A. 110:9583–8. doi:10.1073/pnas.1303483110.

Jiang, F., and J.A. Doudna. 2017. CRISPR–Cas9 Structures and Mechanisms. Annu. Rev. Biophys. 46:505–529. doi:10.1146/annurev-biophys-062215-010822.

Kobayashi, Y., S. Imamura, M. Hanaoka, and K. Tanaka. 2011. A tetrapyrrole-regulated ubiquitin ligase controls algal nuclear DNA replication. Nat. Cell Biol. 13:483–7. doi:10.1038/ncb2203.

Kumar, S., G. Stecher, M. Li, C. Knyaz, and K. Tamura. 2018. MEGA X: Molecular evolutionary genetics analysis across computing platforms. Mol. Biol. Evol. 35:1547–1549. doi:10.1093/molbev/msy096.

Kuroiwa, T., H. Kuroiwa, A. Sakai, H. Takahashi, K. Toda, and R. Itoh. 1998. The division apparatus of plastids and mitochondria. Int. Rev. Cytol. 181:1–41.

Langmead, B., and S.L. Salzberg. 2012. Fast gapped-read alignment with Bowtie 2. Nat. Methods. 9:357–359. doi:10.1038/nmeth.1923.

Lau, O.S., and X.W. Deng. 2012. The photomorphogenic repressors COP1 and DET1: 20 years later. Trends Plant Sci. 17:584–593. doi:10.1016/j.tplants.2012.05.004.

Liao, Y., G.K. Smyth, and W. Shi. 2014. FeatureCounts: An efficient general purpose program for assigning sequence reads to genomic features. Bioinformatics. 30:923–930. doi:10.1093/bioinformatics/btt656.

Markwardt, M.L., G.J. Kremers, C.A. Kraft, K. Ray, P.J.C. Cranfill, K.A. Wilson, R.N. Day, R.M. Wachter, M.W. Davidson, and M.A. Rizzo. 2011. An improved cerulean fluorescent protein with enhanced brightness and reduced reversible photoswitching. PLoS One. 6. doi:10.1371/journal.pone.0017896.

Martin, W., and K. Kowallik. 1999. Annotated english translation of mereschkowsky’s 1905 paper ‘Über natur und ursprung der chromatophoren impflanzenreiche.’ Eur. J. Phycol. 34:287–295. doi:10.1080/09670269910001736342.

Matsuzaki, M., O. Misumi, T. Shin-i, S. Maruyama, M. Takahara, S.-Y. Miyagishima, T. Mori, K. Nishida, F. Yagisawa, K. Nishida, Y. Yoshida, Y. Nishimura, S. Nakao, T. Kobayashi, Y. Momoyama, T. Higashiyama, A. Minoda, M. Sano, H. Nomoto, K. Oishi, H. Hayashi, F. Ohta, S. Nishizaka, S. Haga, S. Miura, T. Morishita, Y. Kabeya, K. Terasawa, Y. Suzuki, Y. Ishii, S. Asakawa, H. Takano, N. Ohta, H. Kuroiwa, K. Tanaka, N. Shimizu, S. Sugano, N. Sato, H. Nozaki, N. Ogasawara, Y. Kohara, and T. Kuroiwa. 2004. Genome sequence of the ultrasmall unicellular red alga Cyanidioschyzon merolae 10D. Nature. 428:653–657. doi:10.1038/nature02398.

Mereschkowsky, C. 1905. Über Natur und Ursprung der Chromatophoren im Pflanzenreiche. Biol. Zent. Bl. 25:593–604.

Minoda, A., R. Sakagami, F. Yagisawa, T. Kuroiwa, and K. Tanaka. 2004. Improvement of culture conditions and evidence for nuclear transformation by homologous recombination in a red alga, Cyanidioschyzon merolae 10D. Plant Cell Physiol. 45:667–671. doi:10.1093/pcp/pch087.

Miyagishima, S., K. Nishida, T. Mori, M. Matsuzaki, T. Higashiyama, H. Kuroiwa, and T. Kuroiwa. 2003. A plant-specific dynamin-related protein forms a ring at the chloroplast division site. Plant Cell Online. 15:655–665. doi:10.1105/tpc.009373.otic.

Nagai, T., K. Ibata, E.S. Park, M. Kubota, K. Mikoshiba, and A. Miyawaki. 2002. A variant of yellow fluorescent protein with fast and efficient maturation for cell-biological applications. Nat. Biotechnol. 20:87–90. doi:10.1038/nbt0102-87.

Naito, Y., K. Hino, H. Bono, and K. Ui-Tei. 2015. CRISPRdirect: Software for designing CRISPR/Cas guide RNA with reduced off-target sites. Bioinformatics. 31:1120–1123. doi:10.1093/bioinformatics/btu743.

Nishida, K., M. Takahara, S. Miyagishima, H. Kuroiwa, M. Matsuzaki, and T. Kuroiwa. 2003. Dynamic recruitment of dynamin for final mitochondrial severance in a primitive red alga. Proc. Natl. Acad. Sci. U. S. A. 100:2146–51. doi:10.1073/pnas.0436886100.

Nishimasu, H., F.A. Ran, P.D. Hsu, S. Konermann, S.I. Shehata, N. Dohmae, R. Ishitani, F. Zhang, and O. Nureki. 2014. Crystal structure of Cas9 in complex with guide RNA and target DNA. Cell. 156:935–949. doi:10.1016/j.cell.2014.02.001.

Nozaki, H., H. Takano, O. Misumi, K. Terasawa, M. Matsuzaki, S. Maruyama, K. Nishida, F. Yagisawa, Y. Yoshida, T. Fujiwara, S. Takio, K. Tamura, S.J. Chung, S. Nakamura, H. Kuroiwa, K. Tanaka, N. Sato, and T. Kuroiwa. 2007. A 100%-complete sequence reveals unusually simple genomic features in the hot-spring red alga Cyanidioschyzon merolae. BMC Biol. 5:28. doi:10.1186/1741-7007-5-28.

Öztürk, N., S.H. Song, S. Özgür, C.P. Selby, L. Morrison, C. Partch, D. Zhong, and A. Sancar. 2007. Structure and function of animal cryptochromes. Cold Spring Harb. Symp. Quant. Biol. 72:119–131. doi:10.1101/sqb.2007.72.015.

Rizzini, L., D.C. Levine, M. Perelis, J. Bass, C.B. Peek, and M. Pagano. 2019. Cryptochromes-Mediated Inhibition of the CRL4Cop1-Complex Assembly Defines an Evolutionary Conserved Signaling Mechanism. Curr. Biol. 29:1954-1962.e4. doi:10.1016/j.cub.2019.04.073.

Sander, J.D., and J.K. Joung. 2014. CRISPR-Cas systems for editing, regulating and targeting genomes. Nat. Biotechnol. 32:347–355. doi:10.1038/nbt.2842.

Suzuki, K., T. Ehara, T. Osafune, H. Kuroiwa, S. Kawano, and T. Kuroiwa. 1994. Behavior of mitochondria, chloroplasts and their nuclei during the mitotic cycle in the ultramicroalga Cyanidioschyzon merolae. Eur. J. Cell Biol. 63:280–8.

Takahara, M., H. Takahashi, S. Matsunaga, S. Miyagishima, H. Takano, A. Sakai, S. Kawano, and T. Kuroiwa. 2000. A putative mitochondrial ftsZ gene is present in the unicellular primitive red alga Cyanidioschyzon merolae. Mol. Gen. Genet. 264:452–460. doi:10.1007/s004380000307.

Takahashi, H., H. Takano, A. Yokoyama, Y. Hara, S. Toh-e, K. Akio, and T. Kuroiwa. 1995. from the primitive red alga Cyanidioschyzon merolae. Curr. Genet. 28:484–490.

Wiese, C., and Y. Zheng. 2006. Microtubule nucleation: γ-tubulin and beyond. J. Cell Sci. 119:4143–4153. doi:10.1242/jcs.03226.

Yagisawa, F., T. Fujiwara, T. Takemura, Y. Kobayashi, N. Sumiya, S.Y. Miyagishima, S. Nakamura, Y. Imoto, O. Misumi, K. Tanaka, H. Kuroiwa, and T. Kuroiwa. 2020. ESCRT Machinery Mediates Cytokinetic Abscission in the Unicellular Red Alga Cyanidioschyzon merolae. Front. Cell Dev. Biol. 8:1–14. doi:10.3389/fcell.2020.00169.

Yoshida, Y., T. Fujiwara, Y. Imoto, M. Yoshida, M. Ohnuma, S. Hirooka, O. Misumi, H. Kuroiwa, S. Kato, S. Matsunaga, and T. Kuroiwa. 2013. The kinesin-like protein TOP promotes Aurora localisation and induces mitochondrial, chloroplast and nuclear division. J. Cell Sci. 126:2392–2400. doi:10.1242/jcs.116798.

Yoshida, Y., H. Kuroiwa, O. Misumi, M. Yoshida, M. Ohnuma, T. Fujiwara, F. Yagisawa, S. Hirooka, Y. Imoto, K. Matsushita, S. Kawano, and T. Kuroiwa. 2010. Chloroplasts divide by contraction of a bundle of nanofilaments consisting of polyglucan. Science. 329:949–53. doi:10.1126/science.1190791.

Yoshida, Y., H. Kuroiwa, T. Shimada, M. Yoshida, M. Ohnuma, T. Fujiwara, Y. Imoto, F. Yagisawa, K. Nishida, S. Hirooka, O. Misumi, Y. Mogi, Y. Akakabe, K. Matsushita, and T. Kuroiwa. 2017. Glycosyltransferase MDR1 assembles a dividing ring for mitochondrial proliferation comprising polyglucan nanofilaments. Proc. Natl. Acad. Sci. 114:13284–13289. doi:10.1073/pnas.1715008114.

Yoshida, Y., and Y. Mogi. 2019. How do plastids and mitochondria divide? Microscopy. 68:45–56. doi:10.1093/jmicro/dfy132.

